# UBTD1 regulates ceramide balance and endolysosomal positioning to coordinate EGFR signaling

**DOI:** 10.1101/2020.05.20.105775

**Authors:** S Torrino, V Tiroille, B Dolfi, M Dufies, C Hinault, L Bonesso, S Dagnino, JP Uhler, M Irondelle, AS Gay, L Fleuriot, D Debayle, S Lacas-Gervais, M Cormont, T Bertero, F Bost, J Gilleron, S Clavel

## Abstract

To adapt in an ever-changing environment, cells must integrate physical and chemical signals and translate them into biological meaningful information through complex signaling pathways. By combining lipidomic and proteomic approaches with functional analysis, we have shown that UBTD1 (Ubiquitin domain-containing protein 1) plays a crucial role in both the EGFR (Epidermal Growth Factor Receptor) self-phosphorylation and its lysosomal degradation. On the one hand, by modulating the cellular level of ceramides through ASAH1 (N-Acylsphingosine Amidohydrolase 1) ubiquitination, UBTD1 controls the ligand-independent phosphorylation of EGFR. On the other hand, UBTD1, *via* the ubiquitination of SQSTM1/p62 (Sequestosome 1) and endolysosome positioning, participates in the lysosomal degradation of EGFR. The coordination of these two ubiquitin-dependent processes contributes to the control of the duration of the EGFR signal. Moreover, we showed that UBTD1 depletion exacerbates EGFR signaling and induces cell proliferation emphasizing a *hitherto* unknown function of UBTD1 in EGF-driven cell proliferation.

## Introduction

All living organisms perceive variations in their environment and translate them into intracellular signals *via* signaling pathways. In multicellular organisms, disturbances in this signal transduction mechanism induce inappropriate cell behavior and are associated with a plethora of diseases including cancer.

Cellular signaling can be viewed as a finely tuned “space-time continuum” [1]. Receptors activated by their ligands at the plasma membrane are endocytosed, then moved along endocytic compartments to be routed to the lysosomes for final degradation or recycled back to the cell surface. Thus, the signal delivered to the cell is the sum of the signals emitted by the activated receptor at the plasma membrane and during its intracellular trafficking [2]. The responsiveness of these processes requires fast and accurate control in space and time, which is mainly ensured by post-translational modification (PTM) of proteins. Broadly, several aspects of cell signaling and receptor trafficking are regulated by proteolytic or non-proteolytic ubiquitination [2–6].

Protein ubiquitination is a PTM that results in the covalent attachment of one or more ubiquitin to lysine residues of the substrate [7–9]. Ubiquitin conjugation occurs in a sequential three-step enzymatic process involving E1 (ubiquitin activation), E2 (ubiquitin conjugation), and E3 (ubiquitin ligation). In this hierarchical framework, there are only two E1s, over 30 E2s and hundreds of E3s in human, illustrating the large spectrum of E2s that contrasts with the high specificity of E3s in the recognition of substrates [10, 11]. While the interaction of E3s with their substrates confers specificity to the system, E2s are more versatile and are commonly considered as “ubiquitin carriers” with an auxiliary rather than control role. However, the modulation of the functionality of E2/E3 complexes by scaffold proteins has been poorly investigated and few proteins acting on these complexes have been characterized [11, 12].

Ubiquitin domain-containing protein 1 (UBTD1) is an evolutionarily-conserved protein which interacts, both *in vitro* and *in vivo*, with some E2 and E3 enzymes of the ubiquitin-proteasome system (UPS) [13]. Recently, we have shown that UBTD1 controls the degradation of the transcriptional regulator YAP (yes-associated protein) by modulating its ubiquitination [14]. Mechanistically, UBTD1 is a component of the ubiquitination complex that allows the E3 ligase β-TRCP (beta-transducin repeats-containing proteins) to interact with YAP. Coherent with this E2/E3 regulatory function, UBTD1 has been reported to stabilize p53 (Tumor Protein P53) through ubiquitination and further degradation of MDM2 (Mouse Double Minute 2), the E3 enzyme that degrades p53 [15]. Importantly, a decreased expression of UBTD1 is associated with increased cancer aggressiveness and decreased overall patient survival in colorectal, liver, prostate, and lung cancer [13, 14, 16]. Based on these evidences, we have previously suggested that UBTD1 may be an E2/E3 scaffolding protein acting as a tumor suppressor [14].

To provide novel insights into UBTD1 functions, we here investigated the effect of its knock-down in a cell model by combining lipidomic, proteomic and signaling screening. Through this integrated approach, we have uncovered an important role of UBTD1 in Epidermal Growth Factor Receptor (EGFR) degradation and signaling. Indeed, we showed that UBTD1 participates in the processing of EGFR-positive vesicles and controls late endosome/lysosome positioning through p62/SQSTM1 (Sequestosome 1) ubiquitination. UBTD1 also regulates the ubiquitination of lysosomal ceramidase N-Acylsphingosine Amidohydrolase 1 (ASAH1) and modifies membrane lipid composition to inhibit EGFR auto-phosphorylation. UBTD1 coordinates in time and space the EGFR signaling pathway to avoid persistent and inappropriate signaling leading to uncontrolled cell proliferation.

## Results

### UBTD1 depletion induces EGFR self-phosphorylation by modifying membrane lipid composition through ASAH1 ubiquitination

The molecular function of UBTD1 is still largely unknown and its cellular role remains elusive. To provide insights into its cellular functions, we performed lipidomic, proteomic, and phospho-kinase array screens in UBTD1-depleted cells (DU145). Using a phospho-kinase array, we observed that UBTD1 depletion increased (> 1.5-fold) phosphorylation of JNK1/2/3, GSK3 α/β (S9/21) and EGFR (Y1086) (**Figure 1A - figure supplement 1A**), EGFR phosphorylation presented the highest difference compared to control cells (> 2-fold). To strengthen these observations, we confirmed by western-blot an increase in EGFR phosphorylation (Y1086) at steady state, and we also noticed an increase in the total amount of EGFR in UBTD1-depleted cells (**Figure 1B**). This observation led us to repeat the same experiment under EGF stimulation (**Figure 1C - figure supplement B**). As compared to EGF-treated-control cells, UBTD1 depletion drastically increased (5 to 20-fold) the phosphorylation of PKB/Akt, β-catenin, GSK3 α/β, CREB, ERK 1/2, and EGFR. In UBTD1 depleted cells, the phosphorylation status of EGFR is the most dramatically changed (15 to 20-fold), suggesting a close link between UBTD1 and EGFR. We then evaluated the STAT3 signaling pathway, a well described target of EGFR [17, 18]. As expected, UBTD1 depletion increased STAT3 phosphorylation, nuclear translocation and STAT3 target gene expression: Bcl-2, HIF-2, and MMP-2, -9 (**Figure supplement C-E**). These converging results indicate that UBTD1 depletion increases EGFR phosphorylation and amplifies signaling cascades downstream of EGFR. We next wanted to elucidate the underlying mechanism by which UBTD1 controls EGFR phosphorylation. In UBTD1-depleted cells, EGFR phosphorylation is increased at steady state (**Figure 1A-B**) without any change in EGF secretion (**Figure supplement 1F**). Thus, we performed a phospho-kinase array in serum starved medium to test whether EGFR was activated independently of its ligand (**Figure 1D - figure supplement 1G**) [19]. Although no growth factors were present in the medium, the levels of EGFR phosphorylation and its downstream targets (Akt, ERK, CREB, STATs) were still higher in UBTD1-depleted cells than in control cells. This data led us to consider that UBTD1 depletion induces ligand-independent EGFR phosphorylation.

**Figure 1:**
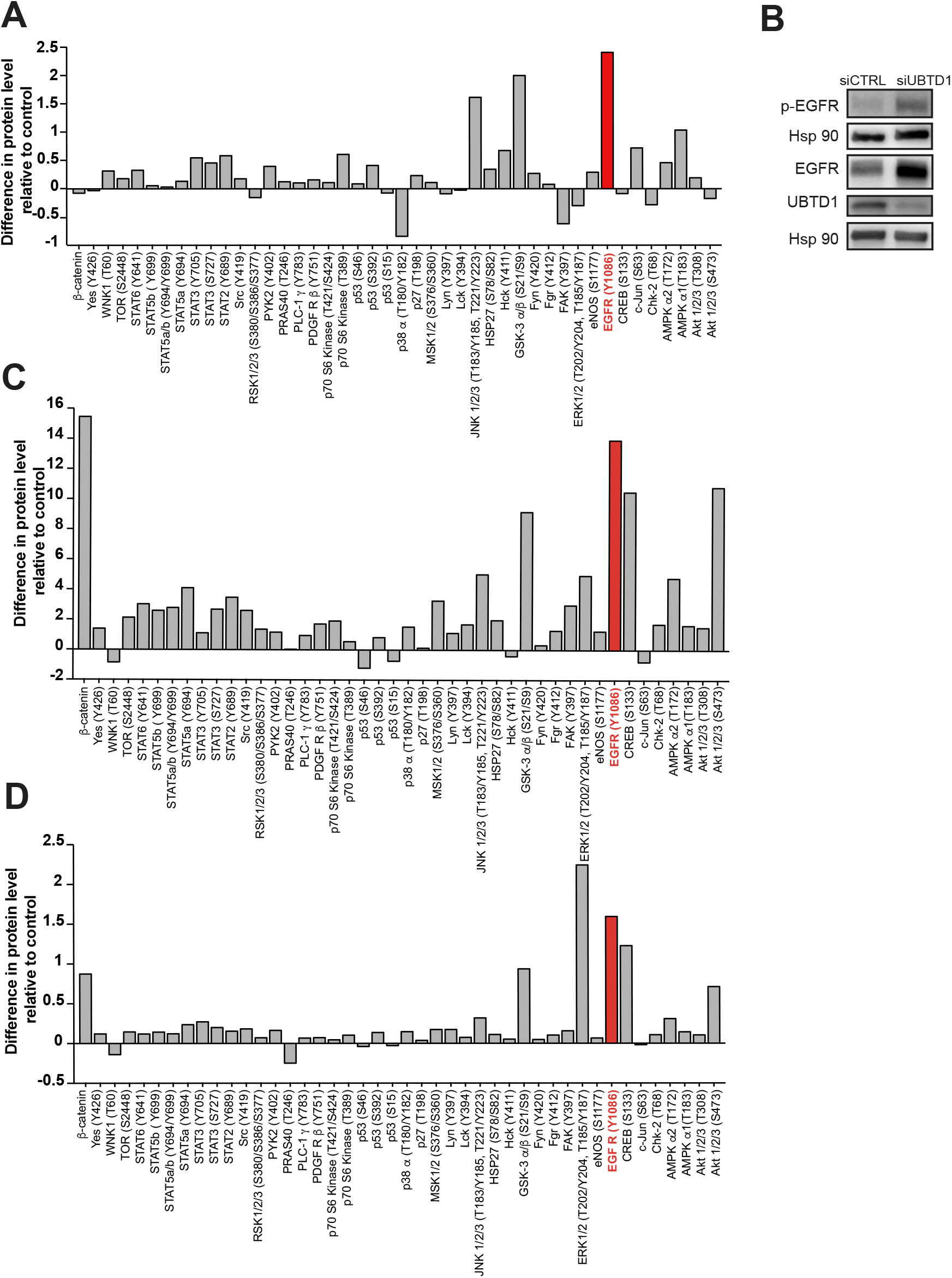
UBTD1 depletion exacerbates EGFR signaling. (A-D) DU145 cells were transfected for 48h with the indicated siRNA (control, siCTRL; UBTD1, siUBTD1). (A,C-D) Relative quantification of protein phosphorylation levels from multiple signaling pathways in complete media (A) or under EGF stimulation (C) or under serum starved medium (D). Data are normalized to the siCTRL condition =0. Phospho-EGFR is in red. (B) Representative immunoblot showing an increase in p-EGFR and EGFR in UBTD1-depleted cells.

Membrane lipid composition surrounding tyrosine kinase receptors can modulate their activation [20–23]. Thus, we isolated cell membranes and performed a lipidomic analysis. UBTD1 depletion induces major changes in membrane lipid composition and, notably, altered the ceramide subclass (**Figure 2A-B**). Next, by MALDI-TOF/TOF mass spectrometry, which detects the non-lipidic moieties and the identity of the ceramide-backboned lipids, we identified a drop in GM2 and GM3 d18:1/16:0 gangliosides (**Figure 2C-D**). The GM3 ganglioside inhibits spontaneous EGFR autophosphorylation [20, 22]. To test whether EGFR phosphorylation induced by UBTD1 depletion was caused by a decrease in GM3 content, we added GM3 to the culture medium. In UBTD1-depleted cells, addition of GM3 restores, in a dose dependent manner, the level of EGFR phosphorylation observed in control cells (**Figure 2E**), suggesting that the increase in EGFR phosphorylation was due to a drop in GM3. To gain insight on how UBTD1 may alter ganglioside level, we performed an immunoprecipitation experiment using endogenous UBTD1 as a bait. Next, by mass spectrometry we identify UBTD1 partners and associated proteins. Using a mild stringency buffer (NP40) to preserve weak protein interactions, we identified a set of 463 proteins (**Source data 1**) distributed in different cell compartments including plasma membrane, endolysosome, and endoplasmic reticulum (ER) (**Source data 2**). Among these proteins, only two are involved in the metabolism of ceramides: ceramide synthase 2 (CerS2) and the lysosomal ceramidase (ASAH1). The protein level of CerS2 was unaffected by UBTD1 invalidation while the level of ASAH1 was significantly increased (**Figure 2F**). In a prostate cancer cell line, it has been suggested that ASAH1 degradation is controlled by the proteasome [24]. To test whether UBTD1 participates in ASAH1 ubiquitination, we performed a ubiquitination assay (**Figure 2G**). UBTD1 knock-down drastically reduced ASAH1 ubiquitination, indicating that UBTD1 is involved in ASAH1 degradation. Hence, we provide compelling evidence showing that UBTD1 depletion induces EGFR phosphorylation by modifying membrane lipid composition through impairment of ASAH1 ubiquitination.

**Figure 2:**
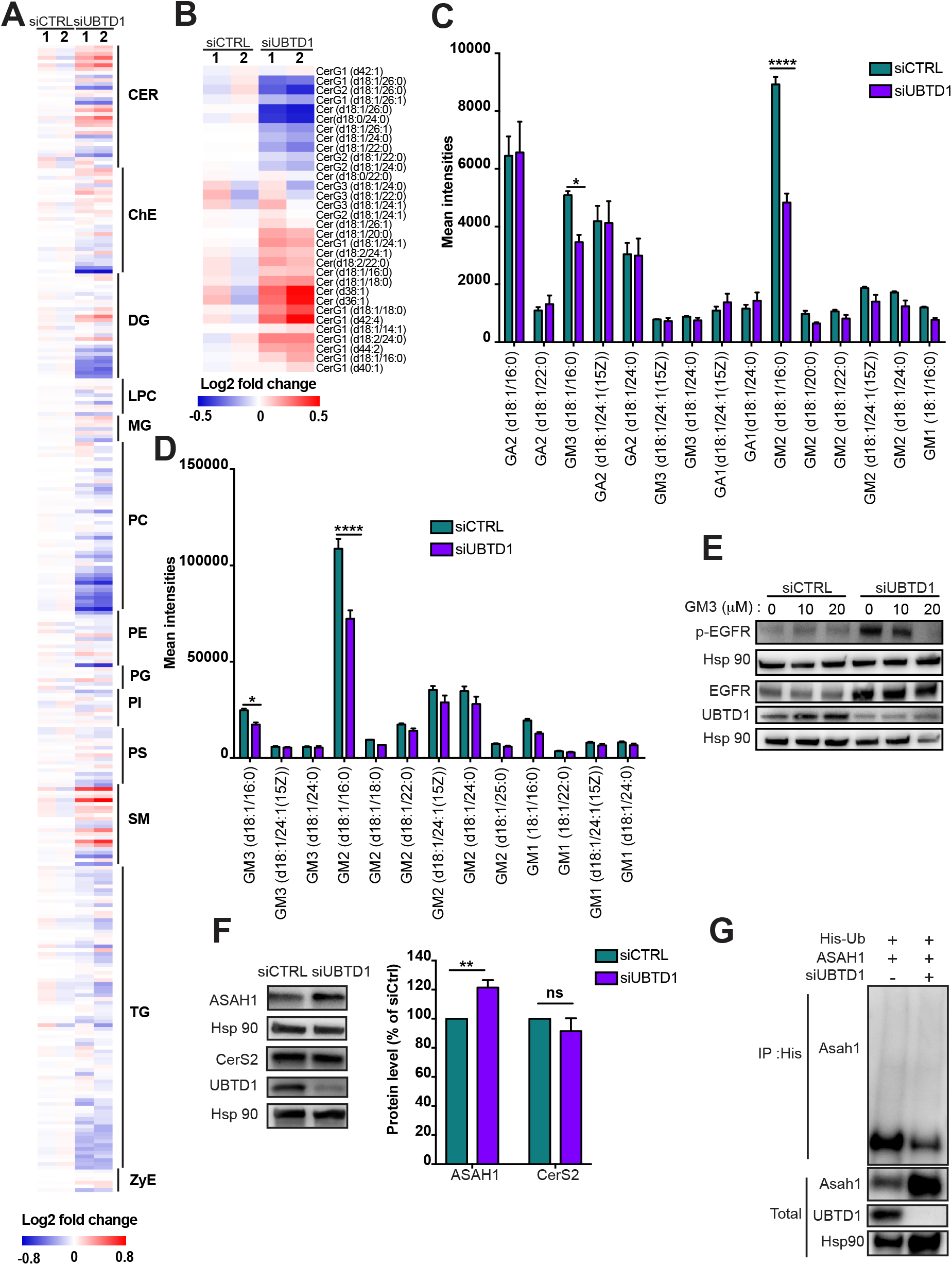
UBTD1 depletion induces EGFR self-phosphorylation by modifying membrane lipid composition through ASAH1 ubiquitination. (A-G) DU145 cells were transfected for 48h with the indicated siRNA (control, siCTRL; UBTD1, siUBTD1). (A-B) Heatmap of all (A) or ceramide (B) lipid levels. The normalized expression of each lipid is shown in a scale range from blue to red. (C,D) Mean intensities of positive-ion (C) and negative-ion (D) MS spectra from reflectron MALDI-TOF analyses. (E) Representative immunoblot of p-EGFR and EGFR in the presence of different concentrations of GM3. (F) Immunoblot and quantification of ceramide synthase 2 (CerS2) and the lysosomal ceramidase (ASAH1). (G) Immunoblots show ASAH1 ubiquitylation in HEK cells in different experimental conditions. Cells were transfected, as indicated, with expression vectors for histidine-tagged ubiquitin (His-Ub) together with control siRNA or UBTD1 siRNA. His-Ub crosslinked forms of ASAH1 were purified (IP: His) and the immunoblot of ASAH1 showed ASAH1 ubiquitylation. The immunoblot of ASAH1 (lower panel) was performed in parallel to verify the amounts of ASAH1 protein engaged in His-Ub purifications. The immunoblot of UBTD1 shows the level of siRNA depletion. n≥3 independent experiments; ns=non-significant; *P<0.05; ****P<0.0001; (C-D, F) two tailed t-test; data are mean ± s.e.m.

### UBTD1 is associated with EGFR and delays its lysosomal degradation

Although the change in the amount of ganglioside GM3 convincingly explains the effect of UBTD1 depletion on EGFR phosphorylation, it does not appear to solely account for the massive effect observed in response to EGF treatment (**Figure 1C**). Indeed, we noticed that the change in EGFR phosphorylation induced by UBTD1 depletion mainly reflects a major increase in total EGFR rather than phosphorylation *per se* (**Figure 1B**), suggesting a predominant role of UBTD1 in EGFR expression or turn-over. To test these two scenarios, we measured EGFR mRNA (**Figure supplement 2A**). No difference in the EGFR mRNA level between control and UBTD1 depleted cells was observed. Thus, we next evaluated the effect of UBTD1 depletion on EGFR turn-over. To decipher the underlying mechanism, we clustered proteins identified in our proteomic experiment based on their STRING interaction scores [25] to sort out the most relevant candidates (**Figure supplement 2B-C**). Consistent with the top-enriched pathway analysis, we discriminated clusters of proteins involved in the ubiquitin-proteasome pathway, RNA processing, cytoskeletal regulation and ER protein processing (**Figure supplement 2D-E**). Interestingly, we identified a cluster centered on EGFR, suggesting that UBTD1 interacts directly with EGFR or with EGFR-associated proteins (**Figure supplement 2C**). Therefore, we next hypothesized that UBTD1 depletion may impair EGFR ubiquitination. To test this possibility, we monitored by proximity ligation assay (PLA) the interaction between ubiquitin and EGFR with or without EGF stimulation (**Figure 3A**). As expected, in control cells, EGF treatment increased EGFR ubiquitination. In UBTD1 knock-down cells, the amount of ubiquitinated EGFR increased similarly to control cells, demonstrating that UBTD1 depletion does not severely impair EGFR ubiquitination. Since the ubiquitination of the EGFR was not modified, we then hypothesized that the intracellular trafficking of EGFR may be impaired. We first determined the amount of EGFR at the cell surface. Using an EGF binding assay, we did not detect any difference between control and UBTD1-depleted cells treated or not with EGF (**Figure 3B**). Next, we performed a time-course of EGF/EGFR endocytosis up to their lysosomal degradation in a pulse-chase experiment using a fluorescent-labeled EGF (**Figure 3C**). The amount of internalized EGF remained constant during the first 30 minutes in both control and UBTD1 knock-down cells, ruling out an internalization defect. In control cells, the amount of EGF started to decay at 120 minutes, whereas in UBTD1-depleted cells degradation was delayed by at least one hour (**Figure 3C**). To determine in which compartment the EGF/EGFR complex was delayed, we combined a pulse-chase of labeled-EGF with immuno-localization of early endosomes (EEA1) or late endosome/lysosome (LAMP1) markers. The arrival and departure of EGF in EEA1-positive compartments was similar in UBTD1-depleted and control cells (**Figure 3D**), suggesting that UBTD1 depletion does not alter the first steps of EGFR intracellular trafficking. However, the extent of colocalization between EGF and LAMP1 strongly increased from the 120 minutes time point in cells depleted for UBTD1 and remained significantly higher for the last two time-points compared to controls (**Figure 3E**). These results underlined that UBTD1 depletion delays the delivery of EGF/EGFR to the degradative lysosomal compartment or impairs the degradative functions of lysosomes.

**Figure 3:**
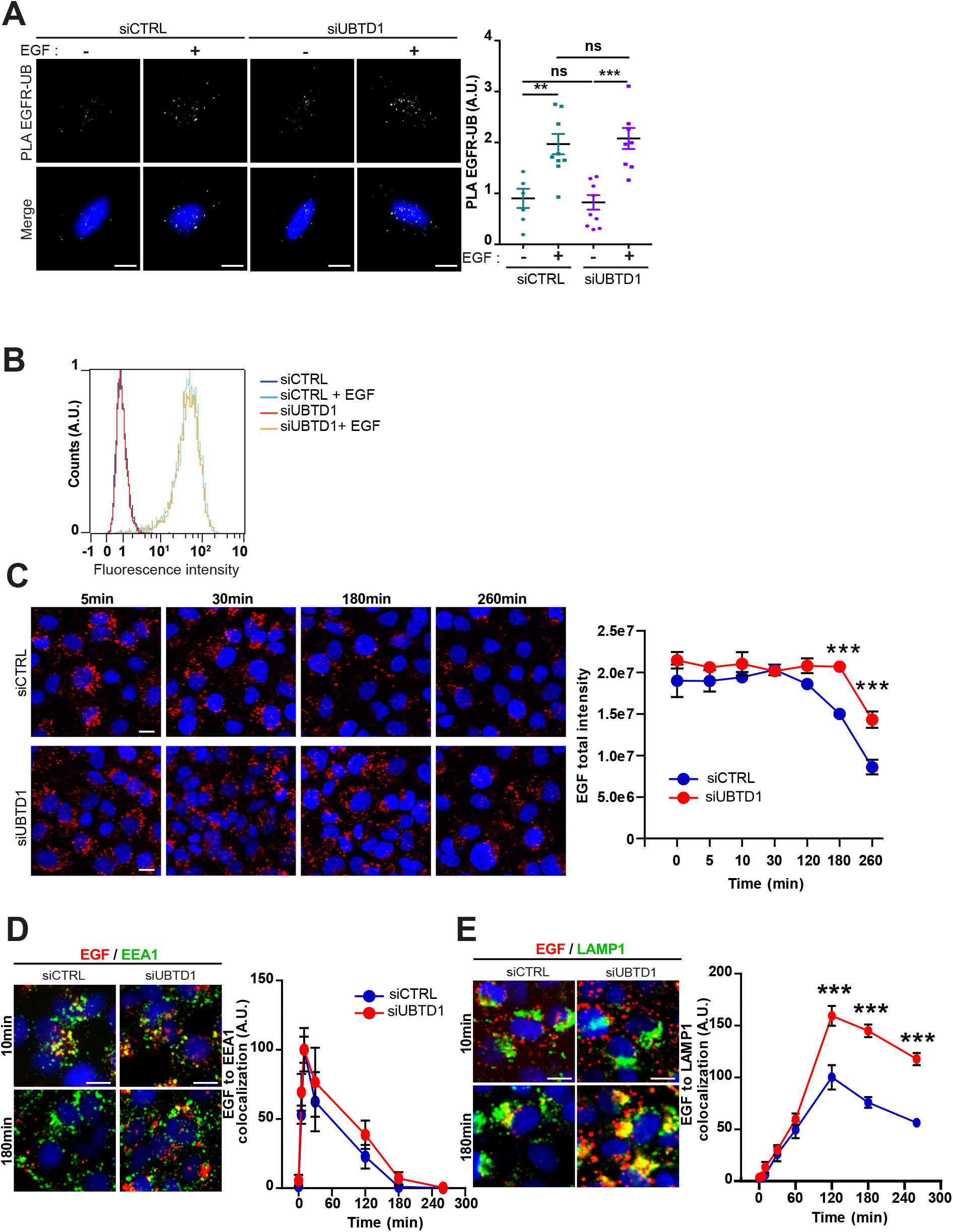
UBTD1 depletion slows-down EGFR degradation. (A-E) DU145 cells were transfected for 48h with the indicated siRNA (control, siCTRL; UBTD1, siUBTD1). (A) Proximal ligation assay monitoring and quantification of EGFR associated with ubiquitin in DU145 treated with EGF. Nuclei were stained with DAPI (blue) on the MERGE image. (B) EGF binding to cell surfaces detected by flow cytometry. (C) EGF-alexa647 pulse chase images and quantification. (D) Representative images and quantification of EGF and EEA1 colocalization during EGF-alexa647 pulse chase. (E) Representative images and quantification of EGF and LAMP1 colocalization during EGF-alexa647 pulse chase. Scale bar=10 μm. n≥3 independent experiments; ns=non-significant; **P<0.01; ***P<0.001; (B, D-F) Bonferroni’s multiple comparison test; data are mean ± s.e.m.

### UBTD1 depletion slows down EGFR degradation without affecting the overall endolysosomal kinetics

We identified many endocytosis-associated proteins as potential UBTD1 interactors in our proteomic analysis (**Source data 1**). Therefore, we first postulated that, in UBTD1-depleted cells, the delay in EGFR degradation could reflect a more general failure of the endolysosomal degradation route. To tackle this question, we analyzed the impact of UBTD1 knock-down on the endocytic compartment’s morphology. During the pulse-chase of EGF, the size and number of EEA1- and LAMP1-positive vesicles was unchanged between control and UBTD1-depleted cells (**Figure supplement 3A-D**). Consistent with this, the number and morphology of the early endosomes, late endosomes, and lysosomes were similar at the ultrastructural level (**Figure supplement 3E-F**). Although the endocytic compartments were present and morphologically normal, the flux of cargoes along these compartments could be altered. To test this possibility, we loaded the cells with quantum dots coupled to BSA (DQ-BSA), a fluid phase cargo that fluoresces when reaching the lysosomes [26]. Both the number of DQ-BSA positive vesicles and the total DQ-BSA intensity were unchanged between control and UBTD1-depleted cells (**Figure supplement 3G-I**). Because the lysosomes were accessible to internalized cargoes and were functional (DQ-BSA fluoresces when lysosomes are functional), it is thus likely that the defect in EGFR degradation observed in UBTD1-depleted cells is not due to a general phenomenon that disrupts the endolysosomal degradation pathway, but is rather restricted to a subset of proteins including EGFR.

### UBTD1 controls p62/SQSTM1 ubiquitination and endolysosomal vesicle positioning

The dynamic of organelles and inter-organelle contacts are essential to allow the exchange of biological materials, but also for spatial coordination of complex cellular processes such as cellular trafficking. The knock-down of UBTD1 does not alter the acidification of the lysosomes as determined with the DQ-BSA nor EGFR trafficking kinetics along the endocytic compartment. Thus, an alternative hypothesis is that UBTD1 affects the spatial positioning of endolysosomal vesicles. To test this proposal, we analyzed the distribution pattern of the early endosome (EEA1) and the late endosome/lysosome (LAMP1) vesicles in control and UBTD1-depleted cells. UBTD1 depletion does not affect the distribution of EEA1-positive vesicles nor their colocalization with the ER marker calreticulin (**Figure 4A - figure supplement 4A**). In contrast, UBTD1 depletion scatters LAMP1-positive vesicles and decreases the colocalization between LAMP1 and calreticulin (**Figure 4B - figure supplement 4B**). Consistent with this, in cells stably over-expressing GFP-tagged UBTD1, the LAMP1-positive vesicles, but not the EEA1-positive vesicles, were more clustered in the perinuclear region (**Figure 4C-D**). To validate this result further, we reproduced this experiment in a human epithelial cell line (RWPE1). Again, UBTD1 over-expression lead to a massive clustering of LAMP1-positive vesicles in the perinuclear region (**Figure 4E**). Taken together, both over- and down-expression experiments support a role for UBTD1 in controlling the positioning of the late endolysosomal degradative compartment.

**Figure 4:**
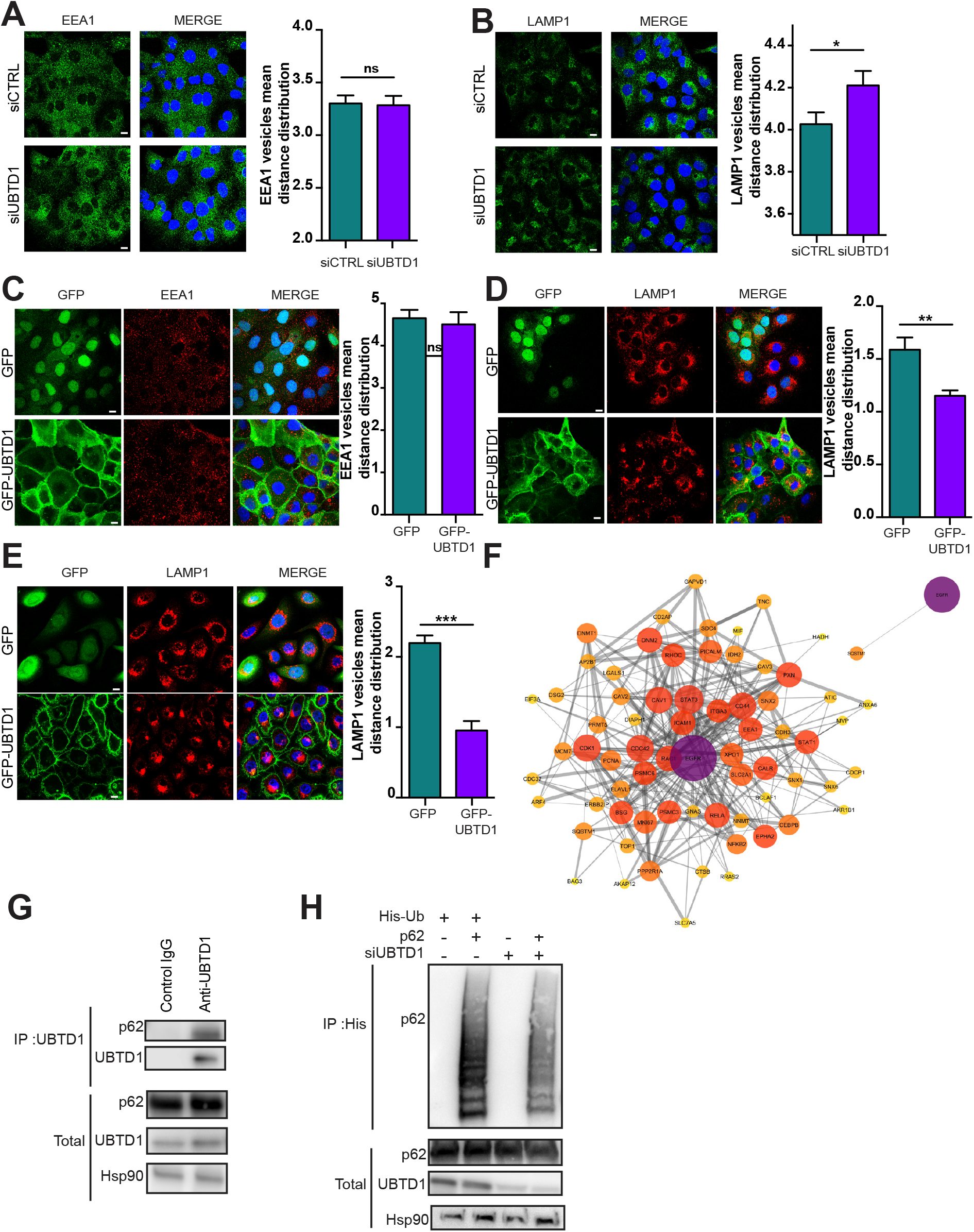
UBTD1 controls p62/SQSTM1 ubiquitination and endolysosomal vesicle positioning. (A-B) DU145 cells were transfected for 48h with the indicated siRNA (control, siCTRL; UBTD1, siUBTD1). Representative confocal immunofluorescence images (left) and quantification (right) of EEA1 (A) or LAMP1 (B) vesicle distribution. (C,D) DU145 or (E) RWPE cells were stably transduced with GFP or GFP-UBTD1. Representative confocal immunofluorescence images (left) and quantification (right) of EEA1 (C) or LAMP1 (D,E) vesicles distribution. (F) UBTD1 partner protein interactome centered on EGFR (first neighbors). Node size (degree: number of connection), edge size (string database combined score). The color scale of the nodes from dark (purple) to light (yellow) reflects the centrality. Minimal protein interaction map derived from left by applying subcellular localization filters (compartment database) focused on endosome, ER and lysosome (StringApp, Cytoscape) (right). (G) Co-immunoprecipitation in DU145 cells between endogenous p62 and UBTD1. UBTD1 was used as bait. The IgG isotype was used as a negative control. (H) Immunoblots show p62 ubiquitylation in HEK cells in different experimental conditions. Cells were transfected, as indicated, with expression vectors for histidine-tagged ubiquitin (His-Ub) together with control siRNA or UBTD1 siRNA. His-Ub crosslinked forms of p62 were purified (IP: His) and the immunoblot of p62 showed p62 ubiquitylation. The immunoblot of p62 (lower panel) was performed in parallel to verify the amounts of p62 protein engaged in His-Ub purifications. The immunoblot of UBTD1 shows the level of siRNA depletion. Scale bar=10 μm. n≥3 independent experiments; ns=non-significant; **P<0.01; ***P<0.001; (A-E) two tailed t-test; data are mean ± s.e.m.

To address how this defect in the positioning of the late endosome/lysosome impacts the degradation of EGFR degradation, we generate a protein-protein interaction network centered around EGFR and applied filters for cell compartment (**Figure 4F**). Using this approach, we identified a minimal interaction network between EGFR and p62/SQSTM1. Next, we confirmed that p62/SQSTM1 co-immunoprecipitated with UBTD1, demonstrating that these two proteins interact or, at least, are in the same protein complex (**Figure 4G**). The ubiquitination of p62/SQSTM1 by the ER-resident E3 ligase RNF26 was shown to organize molecular bridges for the endosolysosomal vesicles to localize in the perinuclear region and is required for efficient cargoes transfer along the endolysosomal degradative route [27].

As depicted in **Figure 4H**, UBTD1 knock-down drastically reduces p62/SQSTM1 ubiquitination suggesting that the correct positioning of EGFR-positive vesicles for their efficient delivery to lysosomes is disrupted by a defect in p62/SQSTM1 ubiquitination.

### UBTD1 depletion triggers an EGFR-dependent cancer cell proliferation

UBTD1 coordinates an EGFR signaling response through the regulation of both the EGFR self-activation and its reduced lysosomal degradation. We next determine how UBTD1 influences cell behavior globally. Consistent with increased EGFR signaling, UBTD1 depletion increased cancer cell proliferation (**Figure 5A-B**). Gefitinib, a FDA-approved EGFR inhibitor, abolished the pro-proliferative effect of UBTD1 knock-down, demonstrating that this effect is EGF-dependent (**Figure 5C**). These results are corroborated by *in silico* data analysis showing that the proteins that we identified using the UBTD1 immunoprecipitation experiment are associated with human carcinoma (**Figure 5D**) and predict a cellular response to gefitinib (**Figure 5E-F**). Despite an absence of correlation at the mRNA levels, UBTD1 mutational status is statistically correlated with EGFR protein levels and cancer patient survival rates (**Figure 5G-I**), partially illustrating the role of UBTD1 on EGFR-post-translational regulation in prostate adenocarcinoma. Although further studies are required, this last observation unveils a plausible link between UBTD1 and EGFR at the post-translational level in humans.

**Figure 5.**
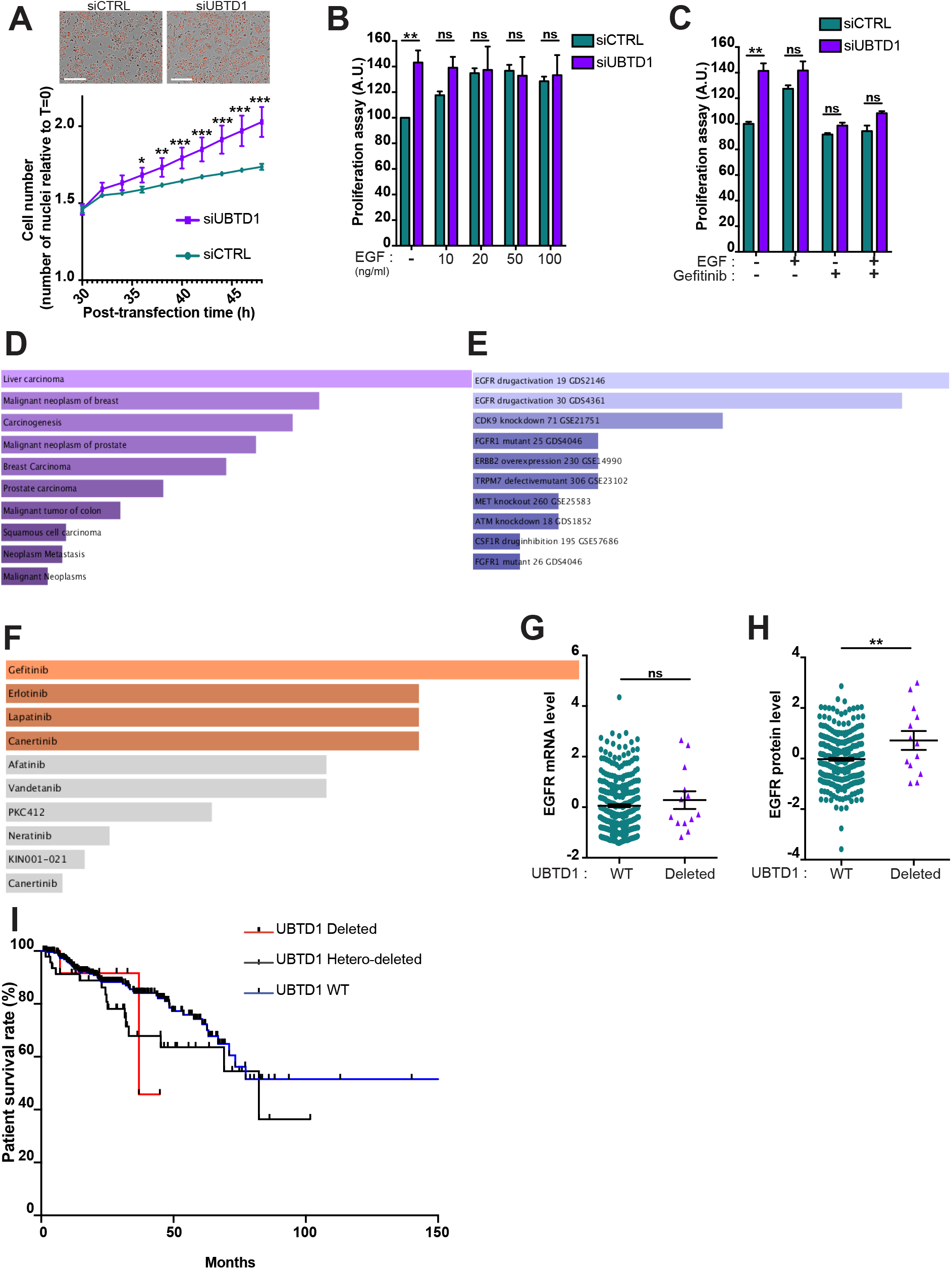
UBTD1 depletion triggers an EGFR-dependent cancer cell proliferation. (A-C) DU145 cells were transfected for 48h with the indicated siRNA (control, siCTRL; UBTD1, siUBTD1). n≥3 independent experiments.(A) Representative images and cell growth curves measured by videomicroscopy (Incucyte, Essen Bioscience). T=0 corresponds to transfection time and the time window (30-50h) is presented. (B, C) Proliferation assay in the presence of different concentrations of EGF (B) and in the presence of EGF inhibitor (Gefitinib) (C). (D-F) Functional interaction analysis was carried out from genes/proteins list identified by mass spectrometry as putative UBTD1 interactants using Dis Genet (D), Kinase Perturbations (E) and KinomScan (F) database. The ranking is established using p-value score. (G) The levels of EGFR mRNA (z-score) in prostate adenocarcinoma patients is compared between patients with or without homo-deletion of UBTD1 gene. (H) The levels of EGFR protein (z-score) in prostate adenocarcinoma patients is compared between patients with or without homo-deletion of UBTD1 gene. (I) DFS were calculated from prostate adenocarcinoma patient subgroups with homo-deletion of UBTD1 gene, hetero-deletion of UBTD1 gene or WT UBTD1 gene.ns=non-significant, *P<0.05 **P<0.01, ***P<0.001; (A-C) two tailed t-test; data are mean ± s.e.m.

Collectively, we here showed that UBTD1 plays a pivotal role in the post-translational regulation of EGFR (**Figure supplement 5A**). By modulating the cellular level of ceramides through ASAH1 ubiquitination, UBTD1 controls the ligand-independent phosphorylation of EGFR. Concomitantly, UBTD1, *via* the ubiquitination of SQSTM1/p62 and positioning of late endosome/lysosome, participates in the lysosomal degradation of EGFR. The coordination of these two ubiquitin-dependent processes contributes to the control of the duration of the EGFR signal and cell proliferation. In conclusion, we have highlighted a yet unsuspected role of UBTD1 on receptor signaling which could be of major importance in certain human pathologies. Moreover, our data lead us to propose that UBTD1 may represent a multistrata coordinator orchestrating EGFR signaling in space and time.

## Discussion

The ubiquitination process works in a hierarchical framework with few E2 enzymes allowing the ubiquitination of many substrates by association with multiple E3 enzymes [11]. Previously, we found that UBTD1 interacts with a subset of E2s to forms stable stoichiometric complexes [13]. Based on these biochemical results, it seems very likely that UBTD1, through its interaction with some E2s, can modify the ubiquitination of many proteins. Illustrating this assumption, we previously demonstrated that UBTD1 controls the ubiquitination of YAP (Yes Associated Protein) and similar findings have been reported for MDM2 (Mouse double minute 2 homolog) [14, 15]. However, these two specific examples presumably reflect only a small portion of the UBTD1 target proteins and, more importantly, of the cellular processes regulated by UBTD1. By using a holistic approach, we here provide evidence that UBTD1 coordinates the EGFR signaling pathway by controlling two distinct ubiquitin-mediated mechanisms. First, UBTD1 controls the self-phosphorylation of EGFR by modulating the ceramide (GM3) balance through the ubiquitination of the acid ceramidase ASAH1. Additionally, UBTD1 controls the lysosomal degradation of EGFR by adjusting the spatial patterning of late endosome/lysosomes and mediating the ubiquitination of p62/SQSTM1. Together, the deregulation of these two molecular checkpoints causes EGFR self-phosphorylation and increases intracellular EGFR lifetime which leads to the persistence of its signaling and induces cell proliferation.

In UBTD1 depleted cells, the membrane lipid composition was altered. Notably the level of gangliosides GM2 and GM3 was most affected. To investigate the underlying mechanism of this lipid switch, we inspected the UBTD1 potential partners and identified ASAH1. ASAH1 is synthesized in the ER as an inactive proenzyme and must be activated through autocleavage to become active in the lysosome [28, 29]. This process is fundamental since ASAH1 is prominently involved in a genetic lysosomal storage disorder in human (Farber’s disease) [30]. However, the regulatory mechanism remains unclear. Interestingly, it has been reported in prostate cancer cells, that ASAH1 is regulated by the proteasome and that the deubiquitinase USP2 modulates protein half-life, clearly suggesting that, ASAH1 is controlled by a ubiquitination-mediated process before reaching the lysosome [24]. Our findings reinforce this proposal by showing that UBTD1 acts on the ubiquitination of ASAH1. Furthermore, ASAH1 has been described by others as a potentially interacting with the E3 ligase HERC2, which we also identified in our proteomic screen as a putative UBTD1 partner [31]. These data lead us to speculate that the lysosomal content of ASAH1 could be regulated by a ubiquitination process mediated by UBTD1 and HERC2. This hypothesis, although interesting, is still speculative and needs to be further explored in the future. Interestingly, Simmons ‘group nicely demonstrated that the ceramide GM3 prevents EGFR self-phosphorylation [20]. In accordance with this work, we observed that UBTD1 depletion, by decreasing cellular GM3 content, increases EGFR self-phosphorylation. In a broader view, we propose that UBTD1 regulates EGFR self-phosphorylation by controlling GM3 content through ASAH1 ubiquitination.

In UBTD1-depleted cells, the decrease in GM3 does not account, by itself, for the dramatic increase in EGFR signaling induced by EGF treatment. Strikingly, we identified EGFR as a potential UBTD1 interactor. Furthermore, the total EGFR level is strongly increased in UBTD1-depleted cells, suggesting a direct connection between UBTD1 and intracellular EGFR trafficking. Once the canonical clathrin-dependent EGFR endocytosis is saturated by excess EGF, EGFR is rapidly routed toward lysosomal degradation through clathrin-independent endocytosis [2, 32]. This critical switch is controlled by EGFR ubiquitination [33, 34]. We and others showed by antibody-based and antibody-free methods that UBTD1 is localized in various cell compartments including the plasma membrane, ER and vesicles [14, 35]. Therefore, we first speculated that UBTD1 was implicated in EGFR cargoes internalization through ubiquitination of the receptor. Surprisingly, the depletion of UBTD1 had no effect either on the ubiquitination of EGFR, or indeed on overall EGFR internalization. This finding indicates that UBTD1 acts on intracellular EGFR processing downstream of the internalization stage and independently of EGFR ubiquitination. In a comprehensive genome wide screen analysis, it has been reported that UBTD1 knock-down increases total EGF vesicle intensity, without major changes in vesicle number, elongation, area or distance to the nucleus [36]. Consistent with this finding, we here demonstrate that UBTD1 depletion drastically increases intracellular fluorescent-labelled EGF lifetime without altering the morphology or the distribution of the early endocytic vesicles. To discriminate at which stage EGF trafficking is corrupted, we analyzed EGF kinetics throughout its endolysosomal trafficking. These functional experiments clearly emphasize that UBTD1 depletion only affects the late endosome/lysosome step of EGF transport while totally preserving the upstream EGF flux. To identify proteins that could be regulated by ubiquination and may interfere with EGFR during this narrow space-time frame, we generated a protein interaction network centered on EGFR using our proteomic-identified proteins. By this method, we uncovered a minimal interactome between EGFR, UBTD1 and P62/SQSTM1. Interestingly, p62/SQSTM1 ubiquitination relies on a UBTD1-target E2s [37]. Thus, we validated the interaction between UBTD1 and SQSTM1/P62 by Co-IP and demonstrated that UBTD1 participates in p62/SQSTM1 ubiquitination. As elegantly demonstrated by Jongsma *et al.*, SQSTM1/P62 is necessary to control EGFR-positive vesicle spatial positioning. Briefly, ubiquitinated p62/SQSTM1 captures specific endolysosomal vesicle adaptors to permit transient ER/endolysosomal contacts which are required for trafficking of some cargoes including EGFR [27]. Hence, we propose that UBTD1 impairs EGFR-positive vesicle positioning through a mechanism close to the one described by Jongsma *et al.* [27]. Albeit a complete description of this process is difficult to demonstrate experimentally, the depletion of UBTD1 leads to a strong reduction in the colocalization between calreticulin and LAMP1, but not EEA1, confirming a defect in endolysosome/ER contact frequency. Additionally, the analysis of LAMP1-positive vesicle distribution shows that UBTD1 depletion or overexpression drastically alters lysosome distribution. Therefore, we could not exclude that UBTD1 depletion affects EGFR degradation through both a defect in EGFR-positive vesicle positioning and an aberrant lysosome distribution *via* a still elusive mechanism. This last hypothesis may be extended to other signaling receptors and clearly needs further clarification.

Finally, EGFR plays a major role in cell proliferation and is frequently overexpressed or hyperactivated in many epithelial cancer cells [38]. Considering the dual role of UBTD1 on EGFR autophosphorylation and lysosomal degradation, we investigated the effect of a decrease in UBTD1 on cell proliferation. Unsurprisingly, our results show that UBTD1 depletion exacerbates EGFR signaling and induces an EGF-dependent cellular proliferation. This last finding underlines the importance of UBTD1 in EGF-driven cell proliferation and may be further investigated in an EGFR hyper-activated state such as cancer. Collectively, we here demonstrated that UBTD1 acts as a coordinator of EGFR signaling, illustrating that the regulation of E2/E3 enzymes of the ubiquitination system by scaffold proteins may represents a critical but still underestimated control layer for coordination of some signaling pathways.

## MATERIALS AND METHODS

### Reagents and antibodies

EGF (10ng/ml) was purchased from PeproTech. Anti-UBTD1 (HPA034825), sodium chloride, DAPI, Gefitinib (SML1657) and 2,5-dihydroxybenzoic acid (DHB) were purchased from Sigma-Aldrich. Anti-HSP90 (sc-13119), anti-RhoGDI (sc-360) and anti-Ubiquitin (sc-8017) were purchased from Santa Cruz Biotechnology. Anti-pSTAT3 (clone D3A7, 9145), anti-STAT3 (clone 124H6, 9139), anti-p-EGF receptor (2234) and anti-EGF receptor (2232) were purchased from Cell Signaling. Anti-EEA1 (610457) and anti-CD107a/LAMP1 (555798) were purchased from BD Biosciences. Anti-SQSTM1/P62 (GTX100685) was purchased from Genetex. Anti-calreticulin-ER Marker (ab2907) was purchased from Abcam. DQ™ Red BSA (D12051) was purchased from Invitrogen. Ganglioside GM3 (860058) was purchased from Avanti Polar Lipids, HRP-conjugated donkey anti-mouse IgG (715-035-150) and HRP-conjugated anti-mouse IgG (711-035-152) were purchased from Jackson ImmunoResearch Laboratories. Epidermal Growth Factor, biotinylated, complexed to Alexa Fluor 647 Streptavidin (Alexa Fluor 647 EGF complex) was purchased from Molecular Probes, Invitrogen (E35351).

### Cell culture

DU145 cells were purchased from the American Type Culture Collection (ATCC). All cells used in this study were within 20 passages after thawing. DU145 cells were cultured (37°C, 5% CO_2_) in Dulbecco’s Modified Eagle’s Medium (DMEM, Gibco) supplemented with 10% fetal bovine serum (Gibco) and Penicillin/Streptomycin (1%, Gibco). The RWPE-1 cell line was obtained from ATCC (CRL-11609). RWPE-1 cells were maintained in KSFM (Life Technologies) supplemented with 5 ng/mL Epidermal Growth Factor (Life Technologies), 50 mg/mL Bovine Pituitary Extract (Life Technologies) and 1% Penicillin-Streptomycin (Life Technologies). The cells were routinely cultured in a humidified atmosphere with 5% CO2 at 37 °C.

### siRNA and plasmid transfection

Transfections were performed with Lipofectamine RNAiMAX according to the manufacturer’s instructions (Invitrogen) using siRNA SMARTpool ON-TARGETplus Human UBTD1 (80019) or ON-TARGETplus Non-Targeting Control siRNAs) (GE Healthcare). Lipofectamine 2000 was used for plasmid transfection according to the manufacturer’s instructions (11668, Invitrogen). Histidine-tagged ubiquitin (pCI-His-hUbi, #31815) and Ha-tagged p62 (HA-p62, #28027) plasmids were purchased from Addgene. DNA constructs corresponding to the mature form of human UBTD1 were subcloned from pEGFP-N1 (Novagen) to a retroviral vector compatible plasmid (PPRIG) [39]. Cells were transduced with an MLV-based retroviral vector and selected by puromycin.

### Human phospho-kinase antibody array

Relative phosphorylation levels of 43 kinases and 2 related proteins were assessed using the Proteome Profiler Human Phospho-Kinase Array Kit (R&D Systems), according to the manufacturer’s instructions. In brief, cell lysates were incubated overnight with nitrocellulose membranes of the Human Phospho-Kinase Array (R&D Systems). Membranes were then washed, incubated with biotinylated detection antibody cocktails, and then incubated with streptavidin-horseradish peroxidase and visualized using ECL (Millipore) and analyzed on Pxi (Syngene). The signal of each capture spot was measured using the ‘Protein Array Analyzer for ImageJ’ and normalized to internal reference controls.

### Cell proliferation assay

48 hours after siRNA transfection, cells were seeded at 5 × 104 cells/well in a 96-well plate and cultured for 24 hours prior to experiment. All conditions were performed in triplicate. Cells transfected with control siRNA served as control. Proliferation was measured using the Cell Proliferation ELISA BrdU kit (Roche Diagnostics GmbH), according to the manufacturer’s instructions. Briefly, cells were labeled with BrdU at a final concentration of 10 μM/well, for 12 h at 37°C. The cells were then denatured with FixDenat solution and incubated for 120 min with 1:100 diluted mouse anti-BrdU conjugated to peroxidase. After two washings (PBS 1X), the substrate solution was added for 25 min and, after this period, the reaction was stopped with 1 M H2SO4 solution. Absorbance was measured within 5 min at 450 nm with a reference wavelength at 690 nm using an ELISA plate reader.

### Kinetic growth assay

Cells were transduced with NucLight Red Lentivirus and selected with puromycin (XXXμg/ml) for 2 weeks (Essen Biosciences). After siRNA transfection cells were allowed to grow (12 well plates) for 48h under a live cell imaging system (Essen Biosciences). The experiments were carried out 3 times independently. Nine images per well (6 wells per condition) were taken every 2 hours for 48h and analyzed with the Incucyte analysis software. The proliferation rate was calculated with the slope between 0-48h.

### Immunofluorescence & pulse-chase

For immuno-fluorescence analysis, the cells were fixed with PBS/PFA 4% for 10 min and permeabilized with PBS/Triton 100X 0.2% for 5 min. After blocking with PBS/BSA 0.2% for 1h, the cells were then incubated with primary antibodies (1/100) at room temperature for 1 h. Secondary antibodies coupled with Alexa-594 and/or Alexa-488 (A-11012, A-11001, Life technologies) were used at 1/500 for 1h. Nuclei were counterstained with DAPI (Sigma-Aldrich). For the pulse-chase experiments, the cells were incubated for 10 min with 100 ng/ml of EGF-alexa647 at 4°C and chased with fresh media for 0, 5, 10, 30, 60, 120, 180 and 240 minutes at 37°C [36]. Then, the cells were washed, fixed and immunolabelled as described above. Images were obtained in a randomized fashion using an LSM510 confocal microscope (Zeiss) and a Nikon A1R confocal. At least 20 images per condition were analyzed using the unbiased multiparametrics MotionTracking software, as previously described (generous gift from Dr Y. Kalaidzidis, Marino Zerial’s lab) [40, 41].

### Electron microscopy

For ultrastructural analysis, cells were fixed in 1.6% glutaraldehyde in 0.1 M phosphate buffer, rinsed in 0.1 M cacodylate buffer, post-fixed for 1h in 1% osmium tetroxide and 1% potassium ferrocyanide in 0.1 M cacodylate buffer to enhance the staining of membranes. Cells were rinsed in distilled water, dehydrated in alcohols and lastly embedded in epoxy resin. Contrasted ultrathin sections (70 nm) were analyzed under a JEOL 1400 transmission electron microscope mounted with a Morada Olympus CCD camera. For quantification, at least 200 structures were counted from at least 20 images per condition acquired randomly.

### Proximity Ligation Assay (PLA)

The PLA kit was purchased from Sigma-Aldrich, and the assay was performed according to the manufacturer’s protocol. Cells were fixed in 4% PFA for 10 min at room temperature, quenched in 50 mM NH_4_Cl for 10 min, permeabilized with 0.2% Triton X-100 (w/v) for 10 min and blocked in PBS/BSA.

Cells were incubated with primary antibodies for 45 min in PBS/BSA. Coverslips were mounted in Fluoromount with DAPI to stain nuclei. PLA signals were visible as fluorescent dots and imaged using an inverted epifluorescence Leica DM 6000B microscope equipped with an HCX PL Apo 63x NA 1.32 oil immersion objective and an EMCCD camera (Photometrics CoolSNAP HQ). Fluorescent dots were quantified using ImageJ. Cells and nuclei were delineated to create masks. After a Max Entropy threshold, the PLA dots were quantified in both masks with the ImageJ Analyze Particles plugin. All counts were divided by the number of cells.

### Flow cytometric analysis

48 hours after siRNA transfection, expression of cell surface EGFR was monitored by flow cytometry on living cells without permeabilization. After washing in ice-cold PBS, cells were immunolabeled on ice for 10 min with Alexa Fluor 647 EGF complex (Molecular Probes, Invitrogen, E35351) in PBS containing 2% BSA. After two further washes in ice-cold PBS containing 2% BSA and one wash in ice-cold PBS, cells were suspended in PBS and analyzed by flow cytometry on a FACS (MACSQuant VYB, Miltenyi Biotec); data were analyzed using the Cell Quest software. Measurements were compared to the isotopic control (APC-conjugated anti-mouse IgG1, Biolegend clone MOPC-21 #BLE400122, 1/100, Ozyme) to determine background and positivity thresholds. Each experiment was repeated at least three times.

### Western blot assays

Cells were lysed in RIPA buffer (Pierce) or directly in Laemmli’s buffer. After denaturation, protein lysates were resolved by SDS-PAGE and transferred onto PVDF membranes (Millipore). Membranes were blocked with 2% BSA in TBS tween20 0.1% and incubated in the presence of the primary and then secondary antibodies. After washing, immunoreactive bands were visualized with ECL (Millipore) and analyzed on Pxi (Syngene).

### Ubiquitylation Assays

HEK293 cells (5 × 10^6^) were transfected with 10 μg of His-tagged ubiquitin WT expression vectors together with p62 and siRNA targeting UBTD1. Ubiquitylated proteins were recovered by His-tag affinity purification on cobalt resin in urea denaturing conditions, as described [42].

### Immunoprecipitation

For endogenous immunoprecipitation, cells were harvested and lysed in NP-40 buffer containing a protease inhibitor cocktail (Thermo Fisher). The lysates (13,200 rpm, 10 min, 4°C) were incubated for 2h with the indicated antibody (IP) at 4°C. Then, 10 μl Dynabeads protein A (Invitrogen) was added to each aliquot for 45 min at 4°C. Beads were washed 3-5 times with NP-40 and eluted by boiling in 2X sample buffer at 95°C for 10 min. The eluted fractions were analyzed by western blot.

### RNA isolation and RT-PCR from cell lines

Total RNA was extracted by TRIzol reagent according to the manufacturer’s instructions (Invitrogen). RNA quantity and quality were determined using NanoDrop™ One Spectrophotometer (Thermo Scientific). One microgram of total RNA was reverse transcribed into cDNA (A3500, Promega). Real-time quantitative PCR was performed using Fast SYBR Green master mix (Applied Biosystems) on a StepOnePlus System (Applied Biosystems). The gene-specific primer sets were used at a final concentration of 1 μM in a 10 μl final volume. RPLP0 mRNA levels were used as an endogenous control to normalize relative expression values of each target gene. The relative expression was calculated by the comparative Ct method. All real-time RT-PCR assays were performed in triplicate with three independent experiments. Primers are provided below. Primers were published in http://pga.mgh.harvard.edu/primerbank/ EGFR (Forward: AGGCACGAGTAACAAGCTCAC; Reverse:ATGAGGACATAACCAGCCACC); BCL2 (Forward: CGCCCTGTGGATGACT; Reverse: GGGCCGTACAGTTCCA); MMP2 (Forward: TGAGCTATGGACCTTGGGAGAA; Reverse: CCATCGGCGTTCCCATAC); MMP9 (Forward: GAACCAATCTCACCGACAGG; Reverse: GCCACCCGAGTGTAACCATA); HIF2 (Forward: TTGCTCTGAAAACGAGTCCGA; Reverse: GGTCACCACGGCAATGAAAC).

### In-Gel digestion

Protein spots were manually excised from the gel and distained by adding 100 μL of H2O/ACN (1/1). After 10 min incubation with vortexing the liquid was discarded. This procedure was repeated 2 times. Gel pieces were then rinsed (15 min) with acetonitrile and dried under vacuum. Each excised spot was reduced with 50 μL of 10 mM dithiothreitol and incubated for 30 min at 56 °C. Alkylation was performed with 15 μL of 55 mM iodoacetamide for 15 min at room temperature in the dark. Gel pieces were washed by adding successively i) 100 μL of H2O/ACN (1/1), repeated 2 times and ii) 100 μL of acetonitrile. Next, gel pieces were reswelled in 60 μL of 50 mM NH_4_HCO_3_ buffer containing 10 ng/μL of trypsin (modified porcine trypsin sequence grade, Promega) incubated for one hour at 4°C. Then the solution was removed and replaced by 60 μL of 50 mM NH_4_HCO_3_ buffer (without trypsin) and incubated overnight at 37°C. Tryptic peptides were isolated by extraction with i) 60 μL of 1% AF (acid formic) in water (10 min at RT) and ii) 60 μL acetonitrile (10 min at RT). Peptide extracts were pooled, concentrated under vacuum and solubilised in 15 μL of aqueous 0.1% formic acid and then injected.

### NanoHPLC-Q-exactive plus analysis

Peptide separation was carried out using a nanoHPLC (ultimate *3000*, Thermo Fisher Scientific). 5 μl of peptides solution was injected and concentrated on a μ-Precolumn Cartridge Acclaim PepMap 100 C18 (i.d. 5 mm, 5 μm, 100 Å, Thermo Fisher Scientific) at a flow rate of 10 μL/min and using solvent containing H2O/ACN/FA 98%/2%/0.1%. Next peptide separation was performed on a 75 μm i.d. × 500 mm (3 μm, 100 Å) Acclaim PepMap 100 C18 column (Thermo Fisher Scientific) at a flow rate of 200 nL/min. Solvent systems were: (A) 100% water, 0.1%FA, (B) 100% acetonitrile, 0.08% FA. The following gradient was used t = 0min 4% B; t = 3 min 4%B; t = 170min, 35% B; t = 172 min, 90% B; t = 180 min 90% B (temperature was regulated at 35°C). The nanoHPLC was coupled via a nanoelectrospray ionization source to the Hybrid Quadrupole-Orbitrap High Resolution Mass Spectrometer (Thermo Fisher Scientific). MS spectra were acquired at a resolution of 70 000 (200 m/z) in a mass range of 150−1800 m/z with an AGC target 5e5 value of and a maximum injection time of 50ms. The 10 most intense precursor ions were selected and isolated with a window of 2m/z and fragmented by HCD (Higher energy C-Trap Dissociation) with a normalized collision energy (NCE) of 27. MS/MS spectra were acquired in the ion trap with an AGC target 2e5 value, the resolution was set at 17 500 at 200 m/z combined with an injection time of 100 ms. Data were reprocessed using Proteome Discoverer 2.2 equipped with Sequest HT. Files were searched against the Swissprot homo sapiens FASTA database update on September 2018. A mass accuracy of ± 10 ppm was used for precursor ions and 0.02 Da for product ions. Enzyme specificity was fixed to trypsin with two missed cleavages allowed. Because of previous chemical modification, carbamidomethylation of cysteines was set as a fixed modification and only oxidation of methionine was considered as dynamic modification. Reverses decoy databases were included for all searches to estimate false discovery rates and filtered using the Percolator algorithm with a 1% FDR.

### Protein-Protein and functional Interactions analysis

protein-protein interaction map and protein clustering were done using Cytoscape software, MCL cluster tool and the String database [25]. Enrichment analysis were performed using Enrichr [43].

### Lipid extraction and analysis

Extraction was performed using 1.5 mL solvent-resistent plastic Eppendorf tubes and 5 mL glass hemolyse tubes to avoid contamination. Methanol, chloroform and water were each cooled down on wet ice before the lipid extraction. Lipids were extracted according to a modified Bligh and Dyer protocol. The cell pellet was collected in a 1.5 mL Eppendorf tube and 200 μL of water was added. After vortexing (30s), the sample was transferred to a glass tube containing 500 μL of methanol and 250 μL of chloroform. The mixture was vortexed for 30s and centrifuged (2500 rpm, 4°C, 10 min). After centrifugation, 300 μL of the organic phase was collected in a new glass tube and dried under a stream of nitrogen. The dried extract was resuspended in 60 μL of methanol/chloroform 1:1 (v/v) and transferred in an injection vial before liquid chromatography and mass spectrometry analysis. Lipid extraction from the membrane purification was carried out with the same protocol in which the solvent volumes used were divided by two. Reverse phase liquid chromatography was selected for separation with a UPLC system (Ultimate 3000, ThermoFisher). Lipid extracts from cells were separated on an Accucore C18 150×2.1, 2.5μm column (ThermoFisher) operated at 400 μl/min flow rate. The injection volume was 3 μL of diluted lipid extract. Eluent solutions were ACN/H2O 50/50 (V/V) containing 10mM ammonium formate and 0.1% formic acid (solvent A) and IPA/ACN/H2O 88/10/2 (V/V) containing 2mM ammonium formate and 0.02% formic acid (solvent B). The step gradient used for elution was : 0 min 35% B, 0.0-4.0 min 35 to 60% B, 4.0-8.0 min 60 to 70% B, 8.0-16.0 min 70 to 85% B, 16.0-25 min 85 to 97% B, 25-25.1 min 97 to 100% B, 25.1-31 min 100% B and finally the column was reconditioned at 35% B for 4 min. The UPLC system was coupled to a Q-exactive orbitrap Mass Spectrometer (thermofisher, CA); equipped with a heated electrospray ionization (HESI) probe. This spectrometer was controlled by the Xcalibur software and was operated in electrospray positive mode. MS spectra were acquired at a resolution of 70 000 (200 m/z) in a mass range of 250−1200 m/z. The 15 most intense precursor ions were selected and isolated with a window of 1 m/z and fragmented by HCD (Higher energy C-Trap Dissociation) with normalized collision energy (NCE) of 25 and 30 eV. MS/MS spectra were acquired with the resolution set at 35 000 at 200 m/z. Data were reprocessed using Lipid Search 4.1.16 (ThermoFisher). In this study, the product search mode was used, and the identification was based on the accurate mass of precursor ions and the MS2 spectral pattern. Mass tolerance for precursors and fragments was set to 5 ppm and 8 ppm respectively. The m-score threshold was selected at 5 and the ID quality filter was fixed at grades A, B and C. [M+H]+, [M+Na]+ and [M+NH4]+ adducts were searched.

### Mass spectrometric analysis for ganglioside

Ganglioside extraction was performed as described with minor modifications [44]. The aqueous upper layers from two extractions (with a mixture of water–methanol–chloroform, W:M:C=2:2:1) were collected. Two volumes of water were added to precipitate polyglycoceramides. The dried pellet was resuspended with methanol/water (M:W=1:1) and analyzed in reflector-positive and -negative modes on MALDI-TOF/TOF mass spectrometer Ultraflex III (Bruker Daltonics, Bremen, Germany) from 700 to 2500 Da (Dalton). External calibration was performed by spotting peptide calibration standard II (BrukerDaltonics). Each sample was spotted in triplicate and mixed with DHB matrix on a steel target plate. All mass spectra were generated by summing 1,000 laser shots for reflectron ion mode, and 1,000 laser shots for the parent mass. Laser power was adjusted between 15 and 30% of its maximal intensity, using a 200-Hz smartbeam laser. MS spectra were acquired in the reflectron ion mode within a mass range from 500 to 2,500 Da. Reflectron ion mode was chosen to obtain high detection sensitivity and resolution. FlexAnalysis version 3.0 and updated 3.4 provided by the manufacturer were applied for data processing. The Human Metabolome Database or HMDB 4.0 (www.hmdb.ca) was used for peak identifications. Heat map and differential intensity analysis with Limma were done using Phantasus (https://artyomovlab.wustl.edu/phantasus/).

### Statistical analysis

All analyses were performed using Prism 6.0 software (GraphPad Inc.). A two-tailed t-test was used if comparing only two conditions. For comparing more than two conditions, one-way ANOVA was used with: Bonferroni’s multiple comparison test or Dunnett’s multiple comparison test (if comparing all conditions to the control condition). Significance of mean comparison is marked on the graphs by asterisks. Error bars denote SEM.

## Acknowledgements

This work was supported by INSERM, the Côte d’Azur University, and by grants from the French National Research Agency (ANR) through the Investments for the program UCA JEDI to S.C. (ANR-15-IDEX-01), the Young Investigator Program to J.G. (ANR18-CE14-0035-01-GILLERON) and the ITMO, Plan Cancer. ST was supported by the “Fondation de France” and T.B. by the French National Research Agency (ANR-18-CE14-0025). We thank the GIS-IBISA multi-sites platform “Microscopie Imagerie Côte d’Azur” (MICA), funded by the “Conseil Départemental 06” support from ITMO Cancer Aviesan (National Alliance for Lifr Science and Health) within the framework of Cancer Plan. FB is a CNRS investigator. We would like to thank Marino Zerial’s lab for giving us access to the MotionTraffic platform for image quantification.

## Author contributions

S.T., J.G., S.C. conceptualized, designed and performed most of the experiments. V.T. generated stable cell lines and performed confocal microscopy experiments. S.D. analyzed proteomic raw data. L.B. and C.H. performed and analyzed mass spectrometry studies. B.D. performed and analyzed live cell imaging experiment. S.L-G. performed electron microscopy experiments and J.G. analyzed and quantified images. M.I. coordinated the imaging facility and generated tools for image analysis. G.A.S. and F.L. performed mass spectrometry for proteomic and lipidomic analysis. S.T., J.G. and S.C. wrote the manuscript. J.H., M.C., T.B., M.D. and F.B. participated actively in data interpretation and critical revision of the manuscript. All authors reviewed, edited and approved the final manuscript.

