## Supplemental Figures for "UBTD1 regulates ceramide balance and endolysosomal positioning to coordinate EGFR signaling"

This PDF file includes:
Supplement figure 1 to 5

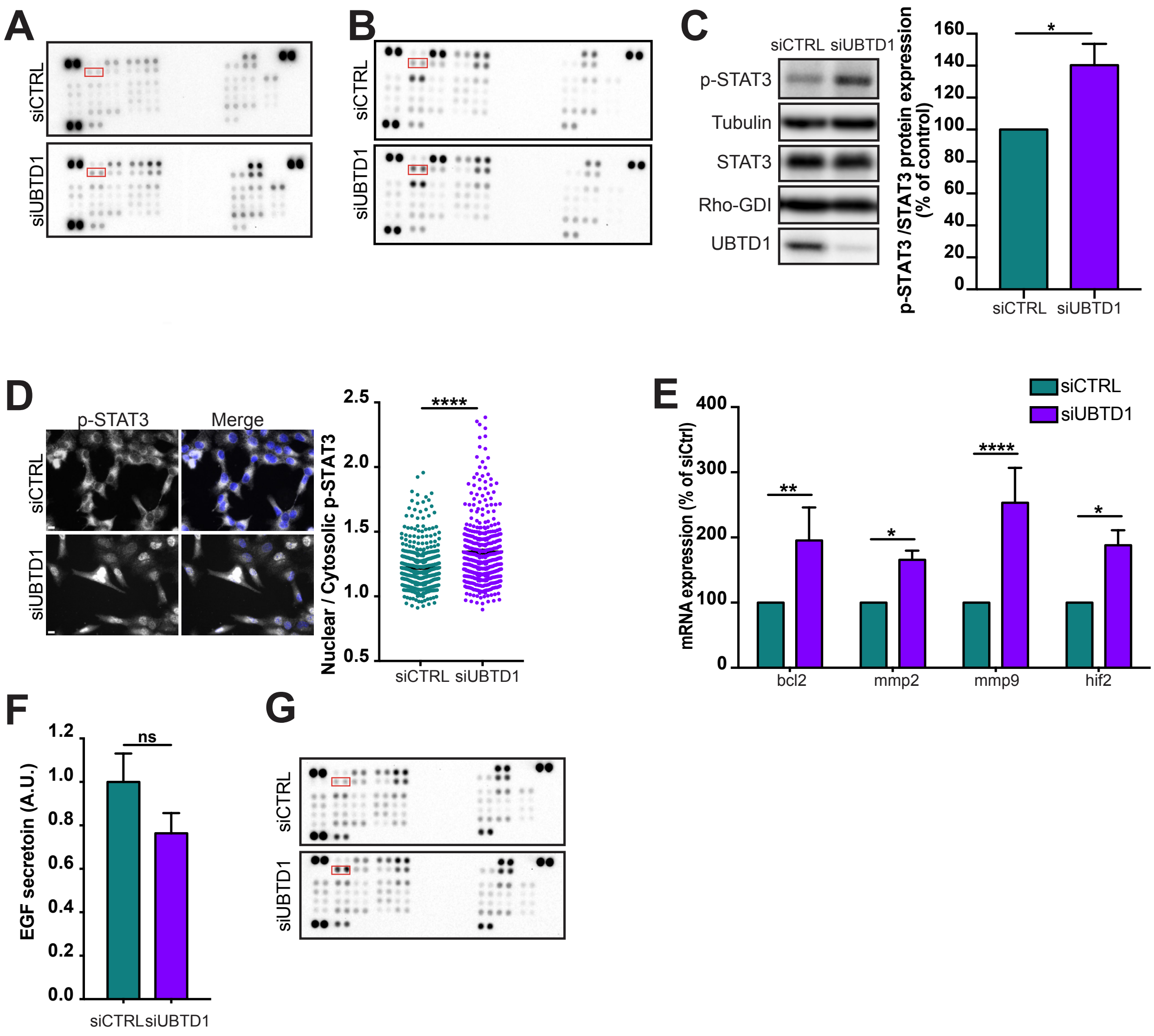


**Supplement figure 1: UBTD1 depletion exacerbates EGFR signaling**

**(A-G)** DU145 cells were transfected for 48h with the indicated siRNA (control, siCTRL; UBTD1, siUBTD1). **(A-B)** Western blotting images of phospho-kinases spotted on the Proteome Profiler Human Phospho-Kinase Array in complete media(A) or under EGF stimulation (B). Phospho-EGFR double spots are marked in red rectangles. **(C)** Immunoblot and quantification (n=3 independent experiments) of p-STAT3. p-STAT3 levels were quantified by calculating the ratio between p-STAT3 and STAT3, both normalized to loading control signal. **(D)** Representative wide-field immunofluorescence images (left) and quantification (right) of pSTAT3 nuclear translocation corresponding to nuclei/cytoplasm mean intensity ratio of pSTAT3. **(E)** mRNA quantification of STAT3 target genes (bcl2, mmp2, mmp9 and hif2). **(F)** EGF secretion measured by ELISA. **(G)** Western blotting images of phospho-kinases spotted on the Proteome Profiler Human Phospho-Kinase Array (serum starved). Phospho-EGFR double spots are marked in red rectangles Scale bar=10 µm. n≥3 independent experiments; ns=non-significant; *P<0.05; **P<0.01; ****P<0.0001; **(C-F)** two tailed t-test; data are mean ± s.e.m.

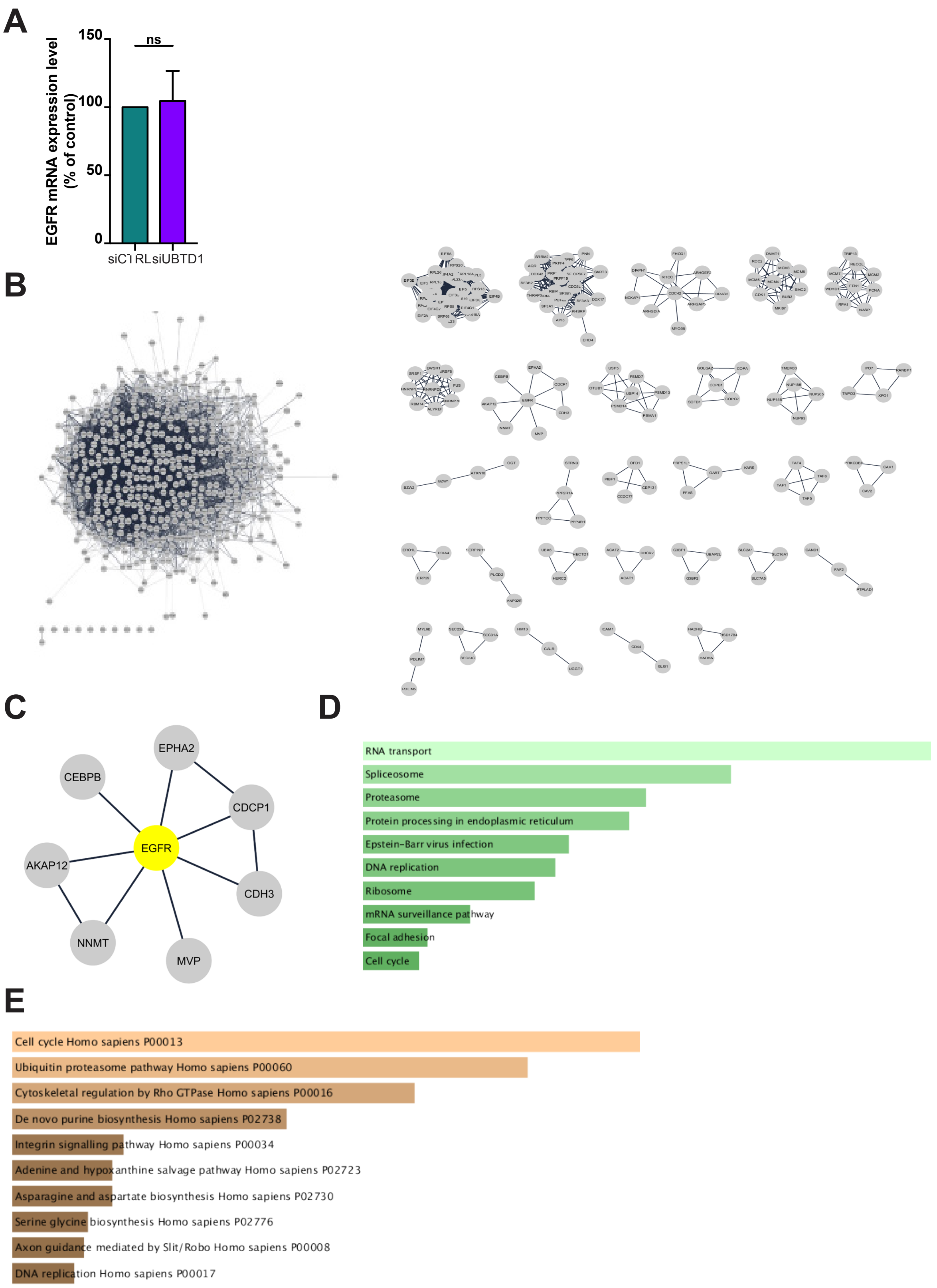


**Supplement figure 2: Protein clustering and functional interaction analysis of the UBTD1 interactome**

(A) Quantification of EGFR mRNA level in DU145 cells transfected for 48h with the indicated siRNA (control, siCTRL; UBTD1, siUBTD1). (B) Clustering of the putative UBTD1 interactants by the Markov Cluster Algorithm method (MCL), using the String database score (inflation 4). (C) Detail of the EGFR centered cluster. (D-E) Functional interaction analysis were carried out from genes/proteins list identified by mass spectrometry as putative UBTD1 interactants using KEGG (D) and Panther (E) database. The ranking is established using p-value score.

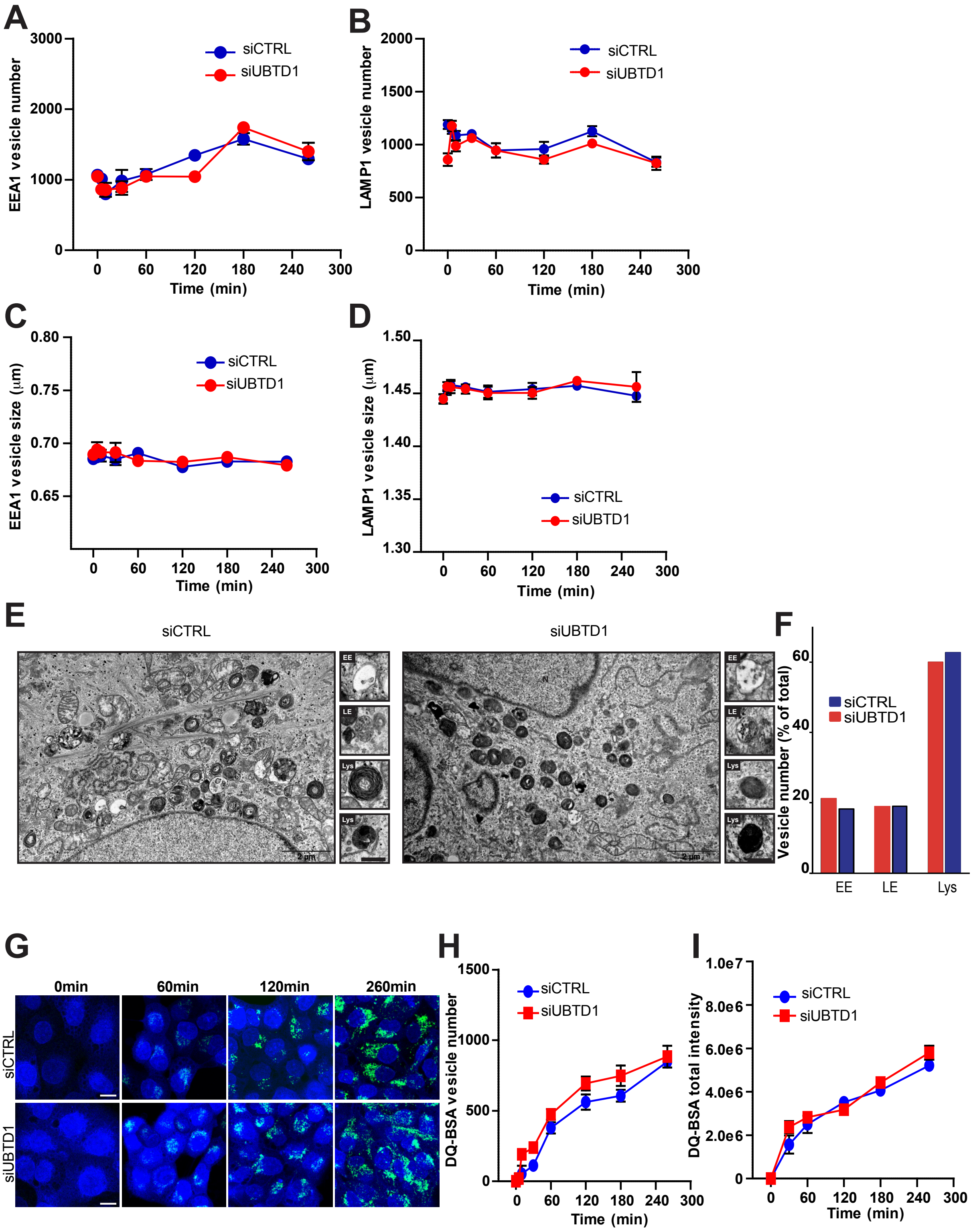


**Supplement figure 3: UBTD1 depletion slows-down EGFR degradation**

**(A-I)** DU145 cells were transfected for 48h with the indicated siRNA (control, siCTRL; UBTD1, siUBTD1). **(A)** Quantification of the EEA1 vesicle number from the pulse-chase experiments. **(B)** Quantification of the LAMP1 vesicle number from the pulse-chase experiments. **(C)** Quantification of the EEA1 vesicle size from the pulse-chase experiments. **(D)** Quantification of the LAMP1 vesicle size from the pulse-chase experiments. **(E,F)** Electron microscopy representative images and quantification of the number of early endosomes, late endosomes and lysosomes based on their morphological description. **(G-I)** Representative images and quantification of the QD-BSA vesicle number and total frame intensity. Scale bar=10 µm. n≥3 independent experiments.

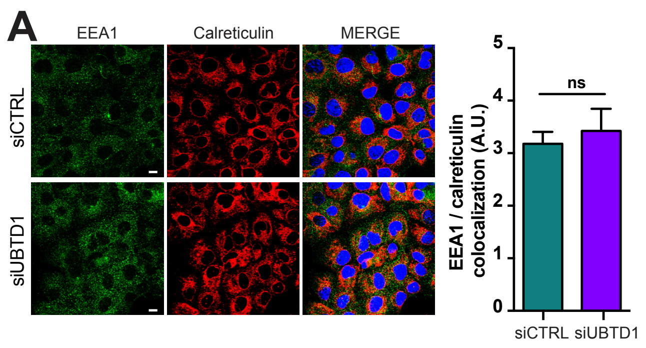


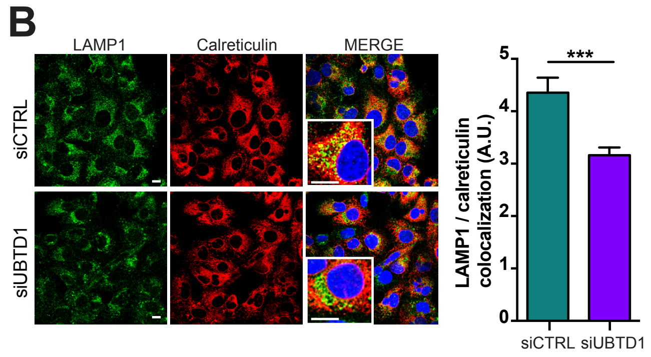


**Supplement figure 4: UBTD1 controls p62/SQSTM1 ubiquitination and endolysosomal v esicles positioning**

**(A-B)** DU145 cells were transfected for 48h with the indicated siRNA (control, siCTRL; UBTD1, siUBTD1). Representative confocal immunofluorescence images (left) and quantification (right) of EEA1 **(A)** or LAMP1 **(B)** colocalization with Calreticulin. Scale bar=10 µm. n≥3 independent experiments; ns=non-significant; ***P<0.001; **(A-B)** two tailed t-test; data are mean ± s.e.m.

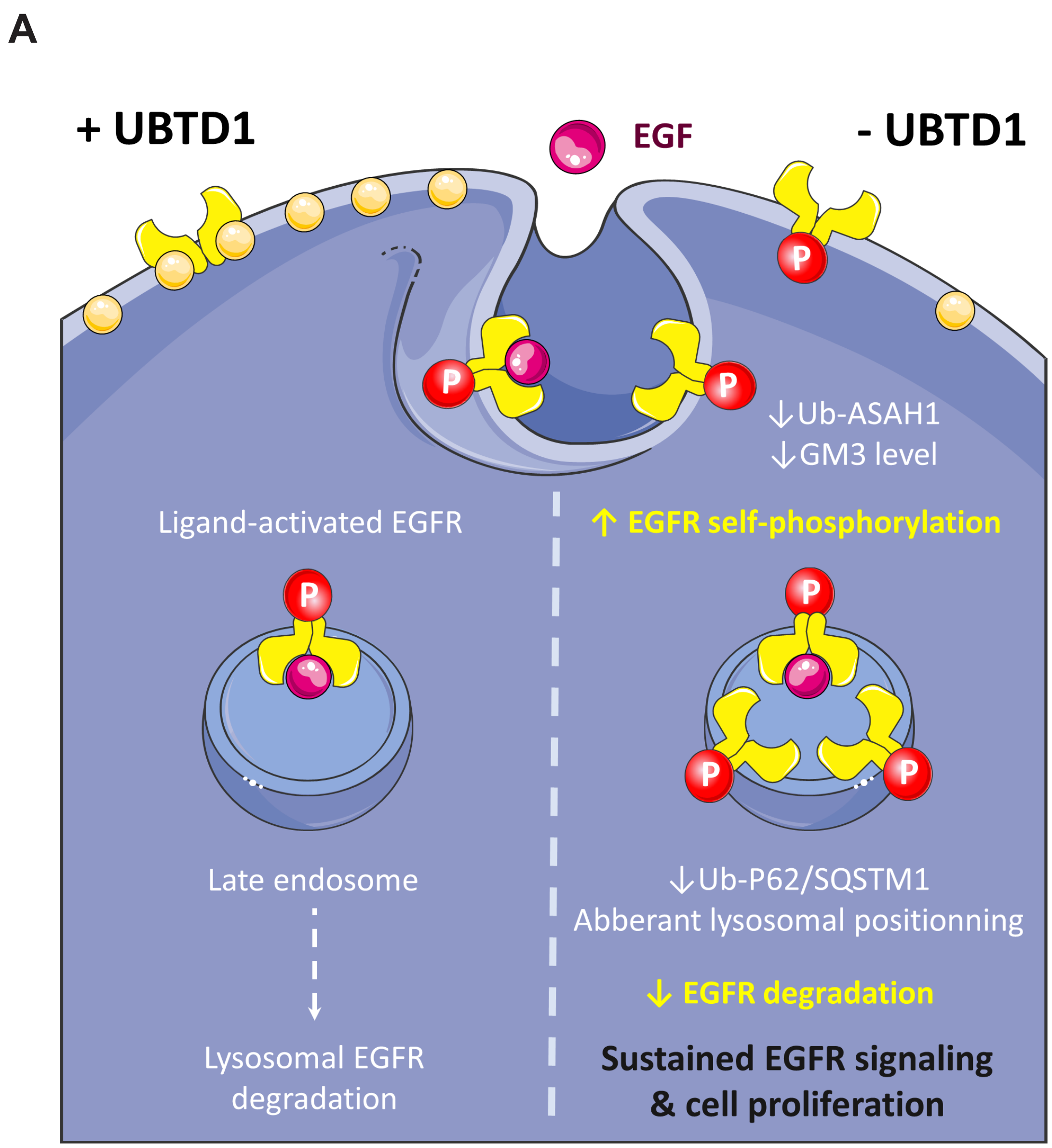


**Supplement figure 5: A proposed model for UBTD1 coordination of EGFR signaling and cell proliferation**

EGF binding induces phosphorylation of EGFR and causes internalization of the EGF/EGFR complex. The complex is then transported by the endocytic system to be either recycled to the membrane or degraded by the lysosome. EGFR auto-phosphorylation is prevented by ganglioside 3 (GM3) (Left). UBTD1 depletion decreases the ubiquitination of the ASAH1 leading to a decline of GM3. Decrease of GM3 induce EGFR self-phosphorylation and signaling. Depletion of UBTD1 also alters endolysosomal positioning and decreases p62/SQSTM1 ubiquitination leading to the inhibition of EGFR degradation. A decrease in EGFR degradation combined with an increase in auto-phosphorylation exacerbates EGFR signaling pathway and induces an EGF-dependent cell proliferation (Right).
